# Structural Determinants of Ligand-Mediated Hmg2 Mallostery

**DOI:** 10.64898/2026.10.30.763229

**Authors:** Christina R. Liem, Jayden Z. Wang, Randolph Y. Hampton

## Abstract

The sterol pathway is a conserved eukaryotic process that produces essential metabolites. Hmg-CoA reductase (HMGR) is a key rate limiting enzyme of the sterol pathway and is the most successful clinical target for managing cholesterol levels. However, determination of the structural features that are essential for its regulation remains elusive. Because of the conservation between mammals and yeast, for many years we have studied the structural features of a yeast Hmg2 isozyme of HMGR. Hmg2 is regulated by the mevalonate pathway isoprenoid GGPP. GGPP acts similarly to an allosteric ligand and appears to bind to a specific site on Hmg2, causing Hmg2 to undergo reversible misfolding. Misfolded Hmg2 is then recognized by the protein quality control mechanism known as the HRD pathway. Due to its similarity to allosteric regulation we have termed this ligand-dependent misfolding “mallostery”. In this work, we systematically probe each residue of the Hmg2 lumenal domain as well as assess the transmembrane domain and lumenal domain as independent domains and use *in vivo* and *in vitro* techniques to assess what regions are important for GGPP response. We identified stabilizing mutations within each loop of the lumenal domain but did not find mutants that were completely unresponsive to GGPP. When assessing the transmembrane domain and lumenal domain as two independent structures, we found that only the transmembrane domain responded to GGPP specifically *in vitro*. Our results suggest that GGPP binding occurs within the Hmg2 transmembrane domain and that the loops in the lumenal domain are required for transducing a mallosteric response to GGPP. Taken together, these data suggest a new model of Hmg2 regulation.

## Introduction

The mevalonate pathway is a conserved eukaryotic biosynthetic route responsible for the production of sterols and a wide range of isoprenoid metabolites essential for membrane integrity, protein modification, and signaling (1, 2). The pathway is largely regulated by the integral endoplasmic reticulum (ER) protein 3-hydroxy-3-methylglutaryl–CoA reductase (HMGR) which catalyzes the reduction of Hmg-CoA to mevalonate. Mammalian HMGR levels are subject to multiple forms of feedback control, including transcriptional, translational, and post-translational regulation (3). Of these, feedback regulated degradation of HMGR is the most selective mode of control. This mode of regulation has been conserved over the billion-year gap between fungi and mammals: in yeast, the Hmg2 isozyme of HMGR undergoes feedback-regulated degradation, providing numerous genetic and biochemical opportunities to study the molecular mechanisms of metabolite-regulated control of enzyme stability (4).

Hmg2 degradation occurs through the Hmg-CoA Reductase Degradation (HRD) branch of the ER-associated degradation (ERAD) pathway which is a conserved quality control system that identifies and eliminates misfolded proteins from the ER (5–7). When sterol pathway flux is high, Hmg2 adopts a conformational state that is recognized by the HRD machinery, leading to its ubiquitination and proteasomal degradation (8–10). Together, this regulatory mechanism and enzyme abundance exemplify the adaptive use of quality control pathways for metabolic feedback regulation.

Our lab has conducted an extensive series of studies to define the mechanism by which metabolic signals control Hmg2 stability. We have previously identified geranylgeranyl pyrophosphate (GGPP), a 20-carbon isoprenoid intermediate of the mevalonate pathway, as a specific and physiologically relevant signal that promotes Hmg2 degradation (11). Our model is that GGPP acts as a high-affinity ligand that induces reversible misfolding of Hmg2, targeting it for HRD-dependent turnover. This metabolite-driven conformational change resembles allosteric regulation, prompting us to describe the mechanism as “mallostery”—ligand-induced protein misfolding to cause quality control degradation (12). GGPP-mediated mallostery is conceptually analogous to monovalent degraders which are small molecules that promote target protein degradation without the need for bifunctional recruitment of degradation machinery (13). Consistent with this model, GGPP-mediated regulation is characterized by high potency, structural specificity, and the presence of antagonistic analogs. Additionally, the requirement for a multimeric structure is consistent with its mallosteric mechanism. Interestingly, the mallosteric response to GGPP requires the conserved sterol-sensing domain (SSD) of Hmg2, a motif present in numerous sterol-regulated proteins, suggesting that SSDs function as general modulators of metabolite-dependent protein regulation (14).

Together, these findings establish GGPP as a metabolic signal that links sterol pathway activity to HMGR turnover through the HRD ERAD pathway. However, despite extensive characterization of GGPP-mediated regulation, the structural basis of this process remains unknown. Specifically, it is unclear where GGPP binds within Hmg2 and whether this interaction occurs at a discrete site or across multiple regions. This gap reflects the broader challenge of defining ligand interactions in multi-pass ER membrane proteins where binding sites are often embedded within the lipid bilayer and difficult to resolve experimentally or computationally (15, 16). For decades, the structural basis of Hmg2 regulation could only be inferred indirectly, through genetics and biochemistry. The advent of accurate structure prediction now makes it possible to test these long-standing hypotheses directly. Here, we combine the AlphaFold-predicted structure of Hmg2 with saturation mutagenesis, domain-specific variants, and *in vivo* and *in vitro* assays to define the structural determinants of GGPP-mediated mallostery.

## Results

### Comparison to Scap, a mammalian SSD protein regulated by sterols, suggested an initial structural hypothesis for Hmg2 mallostery

SREBP cleavage activating protein (Scap) is a mammalian integral ER membrane protein that regulates cholesterol homeostasis by carrying the membrane-anchored transcription factor sterol regulating element-binding protein (SREBP) from the ER to the Golgi. There, SREBP is cleaved and thus activated to transcribe genes encoding sterol pathway enzymes and the LDL receptor (17). Scap protein architecture includes an eight-spanning N-terminal transmembrane domain with an SSD, and a cytosolic C-terminal domain that interacts with SREBP (Fig. 1A).

**Figure 1.**
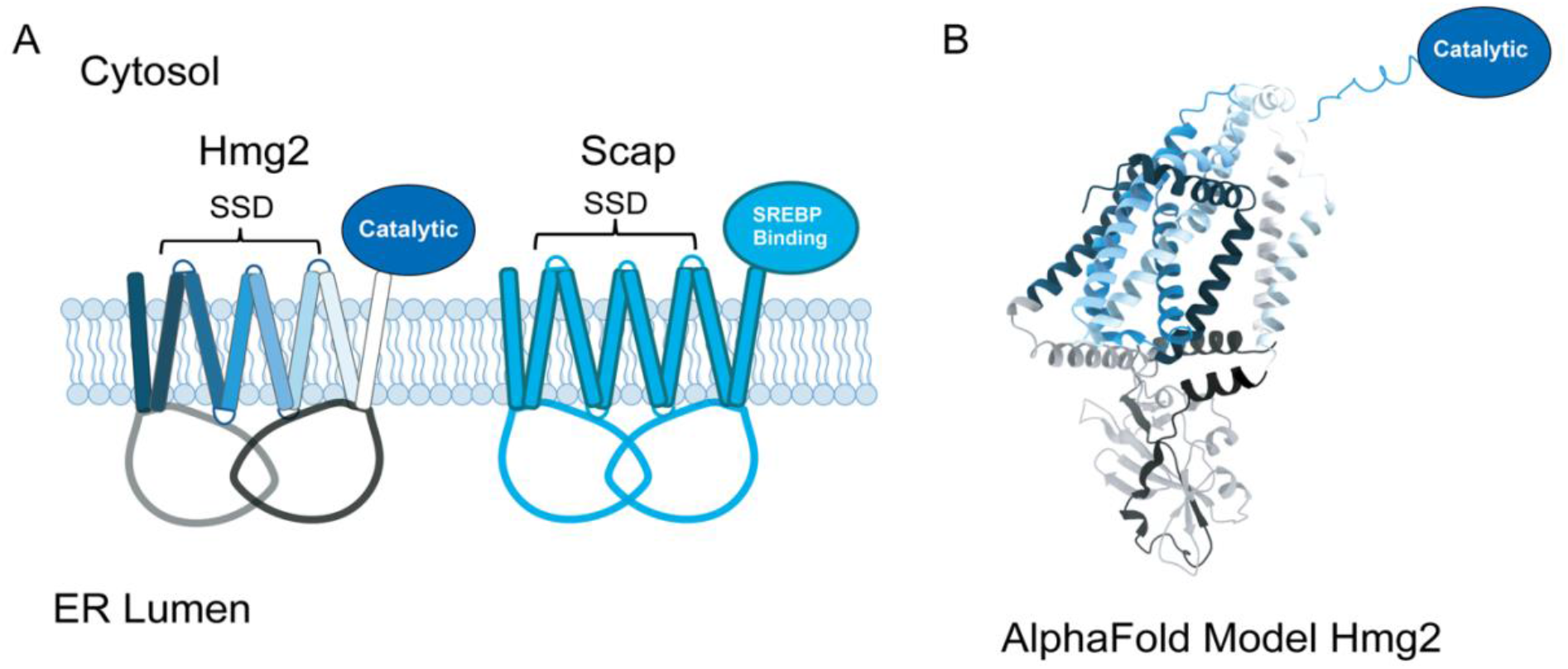
Yeast Hmg2 has structural and functional similarities to mammalian Scap. A, a cartoon model of Hmg2 (left) compared to a cartoon model of Scap (right) embedded in the ER membrane with the SSD highlighted. B, the AlphaFold prediction of Hmg2’s structure shows the eight transmembrane domains with the intertwining of lumenal loop 1 with lumenal loop 7. Domains are colored to match the domains in the cartoon model.

A variety of studies have implicated, somewhat surprisingly, the lumenal loops as being integral for Scap’s binding and response to cholesterol (18–20). Recent cryo-electron microscopy (cryo-EM) studies of Scap reveal that its lumenal loops 1 and 7 exist in an equilibrium between two conformations: (1) directly under the 2-6 transmembrane domain that contains the SSD and (2) angled away from the SSD that allows for more contact with cholesterol embedded in the membrane (21). These data suggest that the position of loop 1 and 7 relative to the SSD modulates Scap activity for regulating cholesterol. In further support of this model, expression of lumenal loop 1 alone is sufficient for cholesterol binding (18), but loop 1 must associate with loop 7 to support functional regulation (19).

Early characterization of Hmg2 established that it is topologically similar to Scap, an integral ER membrane protein with an eight-spanning N-terminal transmembrane domain and a cytosolic C-terminal catalytic domain responsible for mevalonate production from HMG-CoA (Fig. 1A) (22). While this prior work defined several key features of Hmg2 regulation, the GGPP binding site and the structural determinants underlying its response remained unknown. In Scap, its regulatory ligand cholesterol is sensed through two large lumenal loops, loop 1 and loop 7, that intertwine with one another. Based on this precedent, we hypothesized that the analogous lumenal domain of Hmg2 similarly contains the GGPP binding site. Unfortunately, no direct structure of Hmg2 has yet been determined. However, the AlphaFold-predicted structure of Hmg2 reveals two large lumenal loops, loop 1 and loop 7, predicted to intertwine as they do in Scap (Fig. 1B). Accordingly, we posited that mutations of residues involved in GGPP binding or response would stabilize Hmg2 even when GGPP is abundant. Importantly, there is very little sequence homology between Scap and Hmg2; the hypothesis is one of structural analogy more than functional sequence homology. Nevertheless, these striking topological similarities suggested an approach to explore the extent to which the Scap model was functioning in ligand-mediated control of Hmg2 stability.

### A saturated mutagenic screen of Hmg2’s lumenal domain generates phenotypically stable mutants in yeast with high levels of GGPP

To better understand how lumenal loops 1 and 7 domains contribute to GGPP-mediated regulation of Hmg2, we performed a saturated mutagenic screen on these regions to identify all point mutations that alter or disrupt GGPP responsiveness (Fig. 2A).

**Figure 2.**
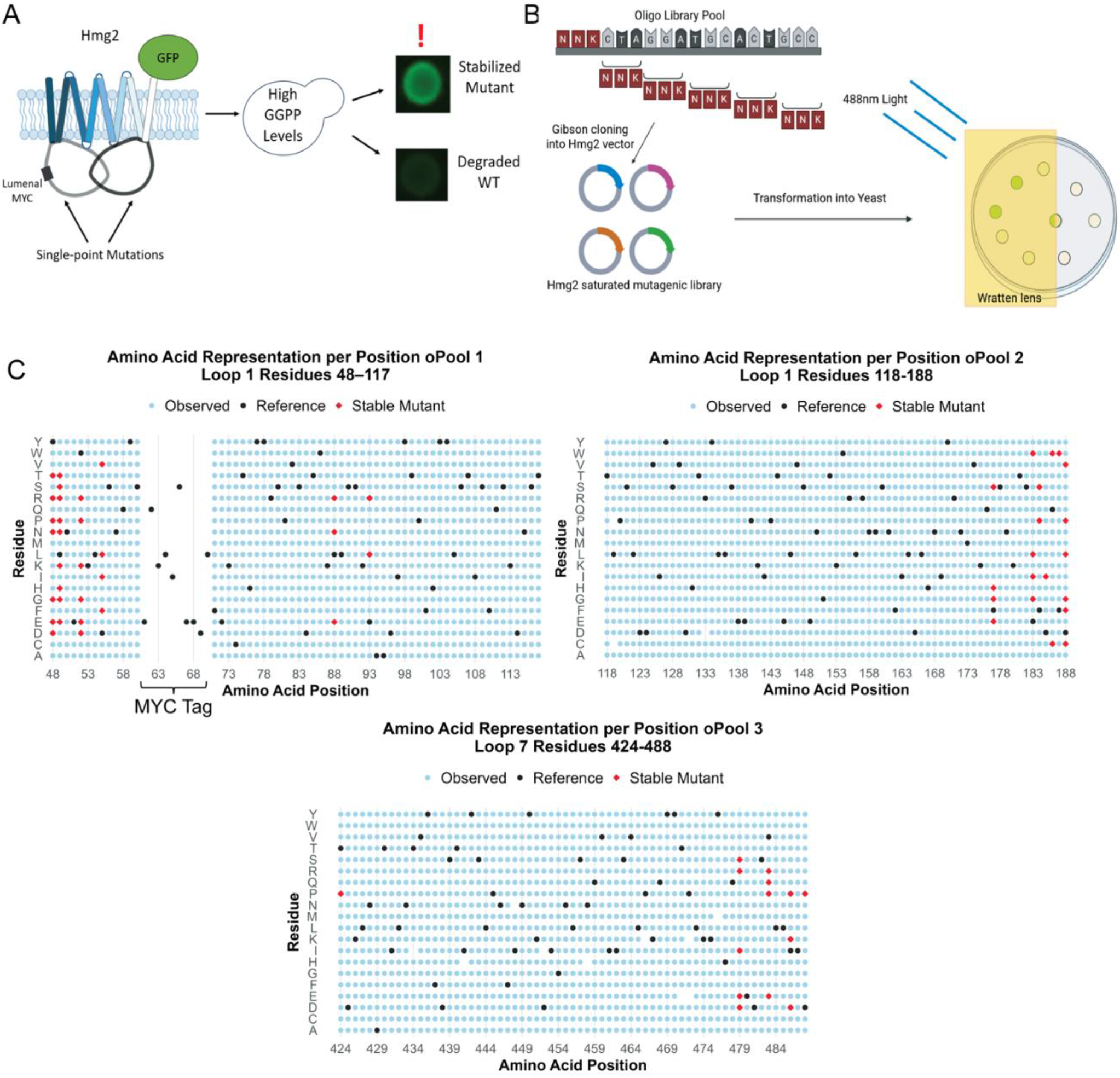
Designing a saturated mutagenic screen of the Hmg2 lumenal domain to find non-responsive mutants. *A,* a schematic of how introducing single point mutations to either loop 1 or loop 7 will be transformed into a yeast strain with high GGPP levels that will produce either stably fluorescing colonies or non-fluorescent colonies. Colonies that were stably fluorescing had the phenotype we were looking for to signify loss of GGPP response and were picked for further analysis. *B*, a schematic of how the saturated mutagenic library was designed. A systematic NNK (N for any nucleotide, K for either a G or T) mutagenesis was introduced at each codon position across either loop 1 or loop 7 to generate all possible amino acid substitutions. These oligos are then cloned into a vector containing the rest of the Hmg2 sequence with Gibson assembly cloning to generate a library of plasmids. These plasmids are then transformed into yeast and screened for fluorescence under a 488nm light source through a wratten lens. *C*, dot plots representing the single amino acid substitutions across lumenal loops 1 and 7 of Hmg2. The x axis represents the position in the library, and the y axis represents each possible amino acid. Black dots represent the WT amino acid, light blue dots represent a single amino acid mutation present in the library, and red dots represent a stably fluorescing mutant identified in the screen. The oPool 1 library has an unmutated region that contains the MYC tag used for subsequent *in vitro* analysis.

We have previously determined that only Hmg2’s transmembrane domain is required for regulated degradation (23). To perform a phenotypic screen, the catalytic domain is replaced with a GFP tag for optical or biochemical detection, and so serves as a proxy for measuring Hmg2 levels and stability both in vivo and by biochemical means. We have also previously shown that placement of a MYC tag in the first lumenal loop does not change Hmg2 regulation and can be used for *in vitro* analysis with a limited proteolysis assay that shows the mallosteric response to GGPP (8, 12, 24–26). Mutants were expressed in a yeast strain with constitutively elevated GGPP levels due to overexpression of the GGPP-synthesizing enzyme Bts1 (11, 12). Under these conditions, wild-type (WT) Hmg2 undergoes rapid regulated degradation and exhibits low fluorescence, whereas mutants that fail to respond to GGPP are stabilized and display increased GFP signal.

Using this design, we hypothesized that mutations disrupting GGPP-mediated mallostery would prevent the conformational transition required for HRD-dependent degradation, resulting in a stable, high-fluorescence phenotype (Fig. 2A). Conversely, mutants that still respond to GGPP undergo HRD-mediated degradation and will appear dim (Fig. 2A). These phenotypic differences can be detected at the colony level with appropriate optics and visual inspection of agar plates.

To systematically identify all such point mutations, saturated mutagenic oligo libraries of loop 1 and loop 7 were generated using an NNK codon strategy to introduce all possible single amino acid substitutions (Fig. 2B). Using Gibson cloning strategies delineated in the Methods, mutant libraries were cloned into a plasmid containing the remaining WT Hmg2 sequence such that each construct encoded a single point mutation. Given the 217 codons targeted, this approach yields ∼4,280 individual variants, which is within the numerical coverage achievable by colony-based screening. Each plasmid library so-generated was deep sequenced to confirm that all amino acids are represented at each position (Fig. 2C).

Following transformation of the resulting mutant library into the yeast screening strain, each colony-bearing plate was screened by illumination with a 488nm light source and examination through a narrow bandpass blue filter to visually detect GFP fluorescence (Fig. 2B). Bright colonies were isolated, re-streaked for purification, and retested for Hmg2-GFP stability using flow cytometry-based assays. Each stable candidate plasmid was isolated from the screening strain and subsequently sequenced individually for mutant identification.

### *In vivo* and *in vitro* analysis of Hmg2 lumenal domain point mutants

We used a combination of *in vivo* and *in vitro* assays to assess how single point stabilizing mutations in the lumenal loops affect Hmg2 stability and GGPP-mediated regulation. Stabilized point mutant-expressing plasmids identified in the screen were isolated and integrated into a wild-type yeast strain with endogenous GGPP production. We first assessed mutant stability using a flow cytometry-based cycloheximide chase assay, comparing each to WT Hmg2-GFP and the mutant Hmg2 S215A-GFP(12, 24, 27).The half-life of WT Hmg2-GFP is about 2 hours in a cycloheximide chase. The S215A mutant renders Hmg2 highly stable due to its unresponsiveness to GGPP and thus shows minimal degradation in a 2-hour cycloheximide chase. Loop 1 and Loop 7 point mutants were analyzed under the same 2-hour cycloheximide chase conditions and compared to these controls (Fig. 3A, Fig. S1A).

**Figure 3.**
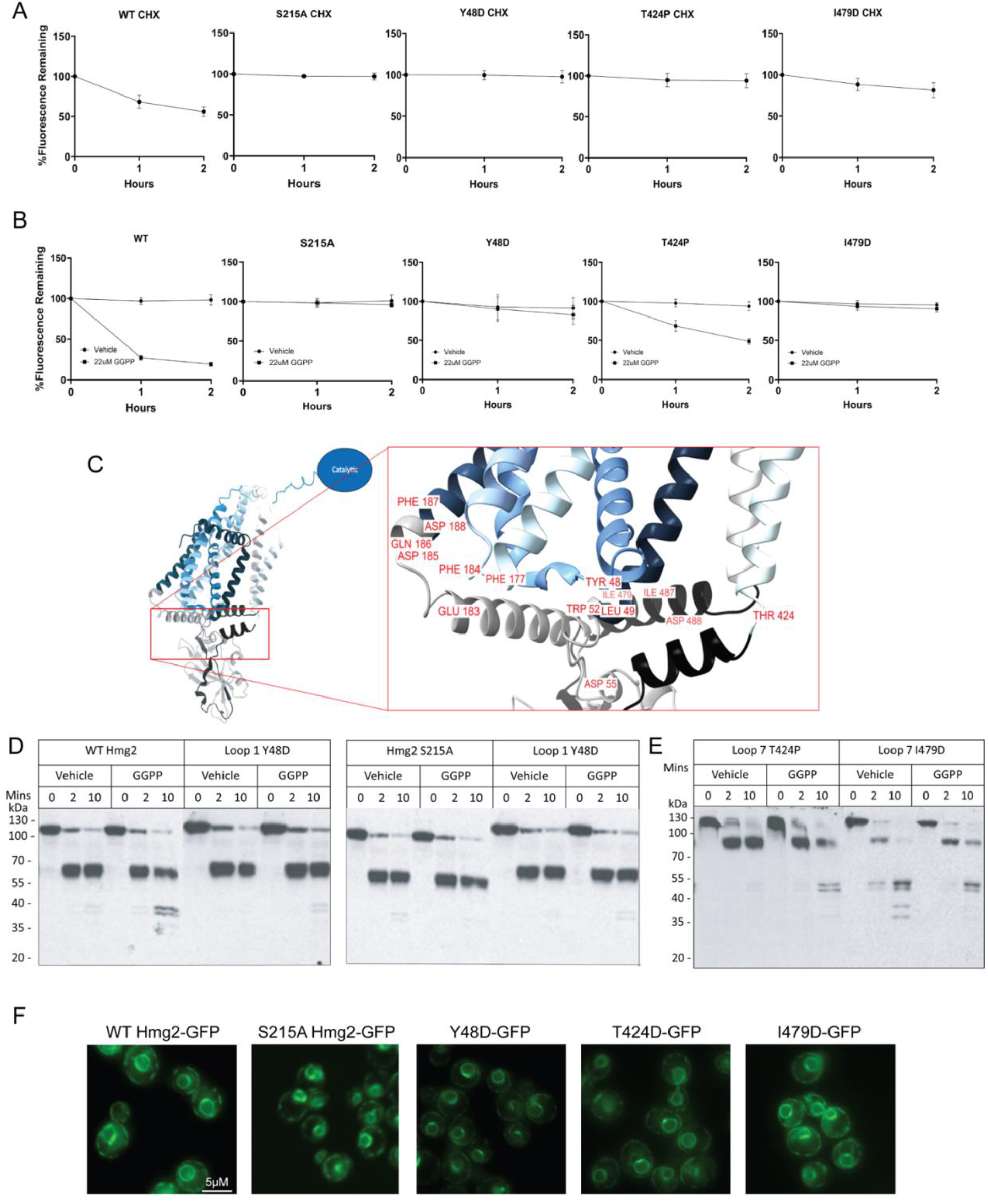
Single point mutations in the lumenal domain have a range of GGPP non-responsiveness compared to WT Hmg2. All flow cytometry experiments show mean fluorescence normalized to time 0, from three separate cultures by flow cytometry of 10,000 cells each. Error bars are SEM. *A*, Loop 1 and Loop 7 mutants were stable compared to the non-responsive SSD mutant S215A. The degradation rate of Hmg2-GFP, SSD mutant S215A, representative loop 1 mutant Y48D, and representative loop 7 mutants T424P and I479D over the indicated times after addition of cycloheximide (CHX). *B*, Loop 1 and Loop 7 have a range of sensitivity to 22uM of GGPP. The effect of GGPP on Hmg2-GFP and Hmg2 mutants S215A, Y48D, T424P, and I479D on steady-state levels over the indicated times. Y48D and I479D are examples of unresponsive mutants like S215A while T424P is an example of a stable mutant with an intermediate response to GGPP. *C*, mapping stable Loop 1 and Loop 7 mutants to the Hmg2 AlphaFold model. All mutants are located relatively close the ER membrane-ER lumen interface. *D*, Loop 1 mutants do not completely block response to GGPP *in vitro*. Limited proteolysis analysis of Y48D compared to WT Hmg2 (left) and S215A (right) over the indicated times. Loop 1 mutants have a blunted response to GGPP *in vitro* when compared to Hmg2 while being more responsive than the stable mutant S215A. *E*, Loop 7 mutants also do not completely block GGPP response *in vitro* and some Loop 7 mutants are stabilized by GGPP *in vitro*. Limited proteolysis analysis of loop 7 mutants T424P (left) and I479D (right). T424P represents loop 7 mutants have the same blunted response *in vitro* as the loop 1 mutants. I479D represents loop 7 mutants that have an inverted response to GGPP and have a decrease in proteolysis when compared to the vehicle. *F*, Representative loop 1 and loop 7 mutants localize to the same subcellular location as WT Hmg2-GFP. Live-cell fluorescence microscopy of WT Hmg2, S215A, Y48D, T424P, I479D. All fluorescent-tagged proteins show a pattern that resembles peripheral ER localization. Scale bar, 5μM.

We next observed mutant Hmg2 responsiveness to GGPP by measuring changes in steady-state fluorescence following the addition of exogenous GGPP with flow cytometry, which gains entry into yeast, allowing direct alteration of the GGPP degradation signal (11, 12, 24). As expected, WT Hmg2-GFP showed a decrease in fluorescence, while S215A remained unchanged. The identified mutants exhibited a range of responses, from WT-like degradation to strongly blunted GGPP sensitivity (Fig. 3B, Fig. S1B). Notably, the majority of these mutants localized to the interface between the lumenal and transmembrane domains (Fig. 3C).

To gain a better understanding of the role of the lumenal loops in the GGPP mallosteric response, a subset of the most strongly affected point mutants were analyzed *in vitro* with our often-employed limited proteolysis assay which provides an empirical readout of Hmg2-GFP’s folding state through its susceptibility to proteolytic cleavage (8, 12, 24). This assay uses microsomes isolated from yeast expressing normally regulated Hmg2-GFP with a single myc tag inserted in the first lumenal loop of the transmembrane domain. Because yeast ER-derived microsomes are oriented with the cytosol facing outward, addition of trypsin exposes the cytosolic face of Hmg2 to proteolysis while the lumenal myc tag remains protected. This tag allows examination of the proteolytic pattern generated by addition of trypsin to the assay, followed by immunoblotting for myc of the resulting fragments. Upon addition of GGPP, Hmg2-GFP limited proteolysis occurs more rapidly without change to the proteolytic pattern itself, while proteolysis of the S215A mutant is unaffected by GGPP. Using this biochemical criterion of GGPP response, we evaluated the most stabilized mutants to gain understanding of the role(s) of the lumenal loops in Hmg2 regulation. Y48D is a representative stable mutant that shows a strongly blunted proteolytic effect when GGPP is added when compared to WT Hmg2 (Fig. 3D, left) and to S215A (Fig. 3D, right). Most stable mutants in either loop 1 or loop 7 isolated by the screen showed a similar lessened response to GGPP in this assay. However, and surprisingly, a small number of loop 7 point mutants showed an “inverted response” to GGPP. Specifically, when GGPP was added, the rate of trypsin-dependent proteolysis was slowed, rather than the normally observed GGPP-dependent increase in rate. I479D is a loop 7 mutant representative of the inverted response mutants, showing a *decrease* in proteolytic rate when GGPP is added (Fig. 3E, right).

To confirm that the point mutants characterized above were suitable for this analysis, we next examined their subcellular localization by fluorescence microscopy. WT Hmg2-GFP, S215A-Hmg2-GFP, Y48D-GFP, T424P-GFP, and I479D-GFP all localized to the cortical ER in a pattern consistent with typical ER membrane insertion (Fig. 3F). These data indicate that the stabilized CHX and GGPP responses observed for these mutants are not attributable to mislocalization or aggregation.

Together, these results demonstrate that lumenal loop point mutations differentially affect Hmg2 stability and GGPP responsiveness. These findings further show that GGPP-induced conformational changes can be altered in both magnitude and direction, indicating that these regions play a critical role in coupling ligand recognition to mallosteric regulation. While the phenotypes of the various mutants clearly indicated that information in the lumenal loops is critical for the GGPP response, they do not resolve the question of whether the loops mediated GGPP binding or response to GGPP bound to a site distinct from loops 1 and 7.

### *in vivo* and *in vitro* analysis of isolated Hmg2 lumenal and transmembrane domains

The above screening results demonstrated the profound importance of the L1 and L7 lumenal loops in the mallosteric response to GGPP. Many single point mutants in either region can drastically alter the *in vivo* response of Hmg2 to GGPP, and these effects can be observed at the level of the structural actions revealed by the limited proteolysis assay. It was further interesting that our collection of point mutants, at least in the AlphaFold model of Hmg2, all appear to be positioned at or near the juxtamembrane region of the Hmg2 molecule. This could indicate significant communication between the lumenal and transmembrane portions as being critical for the mallosteric response. Furthermore, it is clear that the SSD, found within the transmembrane region, also plays a critical role in GGPP-mediated misfolding, since single point mutants like S215A are remarkably stable and unresponsive as well. Accordingly, we decided to ask some broader questions about large scale domains of Hmg2, again informed by the AlphaFold model. To do this, we used an approach inspired by the elegant work of the Radhakrishnan group by examining the L1-L7 domain in isolation by expressing a soluble fusion of the two regions, as well as the Hmg2 transmembrane region free from its lumenal domains. These dramatic alterations help us clarify the roles of these regions of the Hmg2 molecule and revealed a new model of GGPP dependent mallostery.

Specifically, we created two new coding regions to study the importance and possible functions of these Hmg2 domains. A soluble L1-L7-sfGFP fusion was designed by creation of the appropriate coding region, including a signal sequence and a version of GFP that is fluorescent in the ER lumen (Fig. 4A)(28). This L1-L7 fusion was cloned into an integrating yeast expression plasmid, allowing expression and study of this isolated portion of the Hmg2 molecule in the ER lumen. Conversely, we also generated another coding region by study of the AlphaFold model in which the L1 and L7 loops were removed and replaced with small linkers containing a distinct epitope in each replacement loops. This construct was termed “Loopless” due to its lack of either large lumenal loop domain (Fig. 4A). These constructs were integrated into WT yeast strains with normal GGPP production and tested for response in our various *in vivo* and *in vitro* assays.

**Figure 4.**
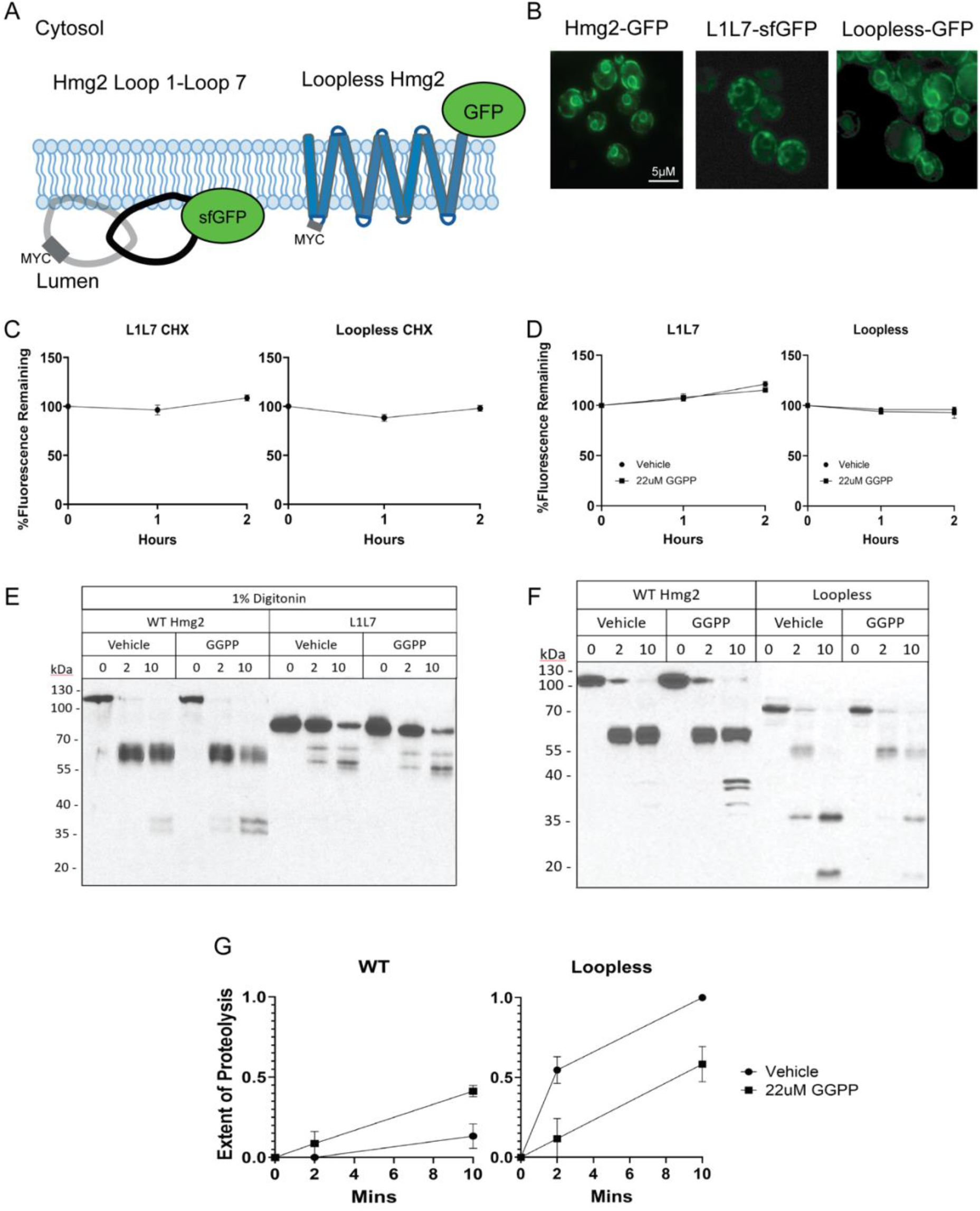
L1L7 and Loopless are stable *in vivo* but Loopless responds *in vitro*. All flow cytometry experiments show mean fluorescence normalized to time 0, from three separate cultures by flow cytometry of 10,000 cells each. Error bars are SEM. *A*, a cartoon depiction of L1L7-sfGFP and Loopless-GFP embedded in the ER membrane with protected 1MYC epitope to allow limited proteolysis assay. *B*, L1L7-sfGFP and Loopless-GFP localize to the same subcellular location as WT Hmg2-GFP. Live-cell fluorescence microscopy of WT Hmg2, L1L7, and Loopless. All fluorescent-tagged proteins show a pattern that resembles peripheral ER localization. Scale bar, 5μM. *C*, L1L7 and Loopless expression remain stable over time *in vivo*. In a cycloheximide chase, the degradation rates of L1L7-sfGFP (left) and Loopless-GFP (right) are measured over the indicated times. *D*, L1L7-sfGFP and Loopless-GFP are unresponsive to 22uM of GGPP. The effect of GGPP on L1L7 (left) and Loopless (right) on steady-state levels are measured over the indicated times. Filled circles show vehicle control and filled squares show GGPP treatment. *E*, digitonin solubilized L1L7-1MYC does not respond to GGPP compared to digitonin solubilized WT Hmg2-1MYC. L1L7 is expressed in the ER lumen and would need to be solubilized for trypsinolysis. WT Hmg2 can be solubilized and trypsinized without changes to its proteolytic pattern. *F*, Loopless-1MYC has a stabilizing response to GGPP when compared to WT Hmg2-1MYC in a limited proteolysis experiment. There is a decrease in rate of proteolysis when Loopless is treated with GGPP. *G*, Densitometry of the proteolytic products in WT Hmg2-1MYC or Loopless-1MYC limited proteolysis show that WT Hmg2 increases in proteolysis while Loopless shows a decrease in proteolysis upon GGPP treatment. Immunoblots were scanned as TIFF files and pixels were counted for total lane intensity. WT Hmg2 Extent of Proteolysis = (Final doublet intensity/Total intensity), p < 0.05. Loopless Extent of Proteolysis = (Doublet intensity/Total intensity), p < 0.05. Values shown are means of three experiments. Error bars are SEM. Statistical significance was determined by two-way repeated-measures ANOVA followed by Sidak’s multiple comparisons test comparing Vehicle and GGPP at each time point.

To confirm that these truncations were relevant for studying Hmg2, we first observed their subcellular localization with fluorescence microscopy. WT Hmg2-GFP shows a characteristic ER pattern (Fig. 4B left). Both the L1L7 and Loopless mutants display an identical pattern of ER fluorescence, indicating that these mutants are localized identically to WT Hmg2 (Fig. 4B middle and right).

Next, we measured their stability with a cycloheximide chase to compare it to WT Hmg2. Surprisingly, and despite the loss of many predicted intraprotein interactions by either modification, both the L1L7 and Loopless mutants were extremely stable, each showing almost no change in fluorescence in a 2-hour CHX chase (Fig. 4C). Furthermore, neither L1L7 nor Loopless showed any change in steady state levels after treatment with exogenous GGPP (Fig. 4D).

Both L1L7 and Loopless were stable and unresponsive to GGPP by *in vivo* optical assays. We next used versions of the limited proteolysis assay to further test if either construct displayed an interaction with GGPP as measured by this method. Because the L1L7 mutant is expressed in the lumen of the ER, microsomes were isolated and detergent-solubilized so that L1L7 could be accessible for trypsinolysis. We have previously shown that full-length Hmg2-GFP retains its *in vitro* response in this assay when so solubilized (24). The multi-spanning Loopless mutant was assayed in an identical manner to Hmg2 above in isolated microsomes without detergent. Our working hypothesis, in analogy to the case with Scap, predicted that the L1L7 construct would interact with GGPP and that Loopless construct would not. Nevertheless, the limited proteolysis assay showed that there were no structural changes *in vitro* when GGPP is added to L1L7 (Fig. 4E). Conversely, and remarkably, Loopless did indeed show a structural response to GGPP: there was a detectable and statistically significant (Fig. 4G, right) decrease in proteolysis when GGPP was present in the assay (Fig. 4F). Specifically, the presence of GGPP caused a slowing of the appearance of the trypsinolysis pattern, indicating a structural effect of – and presumably a binding site for – GGPP. This difference in proteolysis between WT Hmg2 and the Loopless mutant was measured by densitometry to quantify the extent of proteolysis when treated with vehicle and GGPP, respectively. The fraction of WT Hmg2 that gets proteolyzed during a 10 min incubation increases from 0.1 to 0.45 (Fig. 4G left) when treated with GGPP. The fraction of Loopless that gets proteolyzed decreases from 1 to 0.5 when treated with GGPP (Fig. 4G right). This suggests that the Loopless mutant still responds to GGPP but is now being stabilized by its presence. This surprising result indicated, in contrast to our prediction, that Loopless had a binding site for GGPP, while L1L7 appears to not interact, at least by these assays. We next evaluated the features of this Loopless-GGPP interaction, and in particular, we asked how similar this interaction was to that we have extensively characterized in Hmg2 (12, 24, 25).

### Characterizing the Loopless mutant GGPP interaction

We have previously reported that Hmg2 has a specific potent response to GGPP. Perhaps the strongest piece of evidence for a specific, allostery-reminiscent binding site in Hmg2 is the existence of a GGPP-based antagonist: GGSPP (12). This analogue of GGPP shows no mallosteric effect when added to Hmg2 in the limited proteolysis assay, nor any action *in vivo*, when present alone even at high concentrations. However, GGSPP successfully antagonizes the actions of GGPP in both *in vivo* and *in vitro* assays. Furthermore, the very similar analog 2F-GGPP has no antagonistic actions. Thus, in the normal Hmg2, it would appear that GGSPP can bind to its site, while 2F-GGPP cannot. This provides a specific criterion to apply to the GGPP interaction in Loopless. If it has the same binding site, we expect GGSPP to interact with Loopless, while 2F-GGPP should not, as assessed by the limited proteolysis assay. Indeed, Loopless was unresponsive to 2F-GGPP, while GGSPP caused the same structural effect as GGPP (Fig. 5A, B), indicating that the two analogues show the same interaction specificity with Loopless as they do with Hmg2.

**Figure 5.**
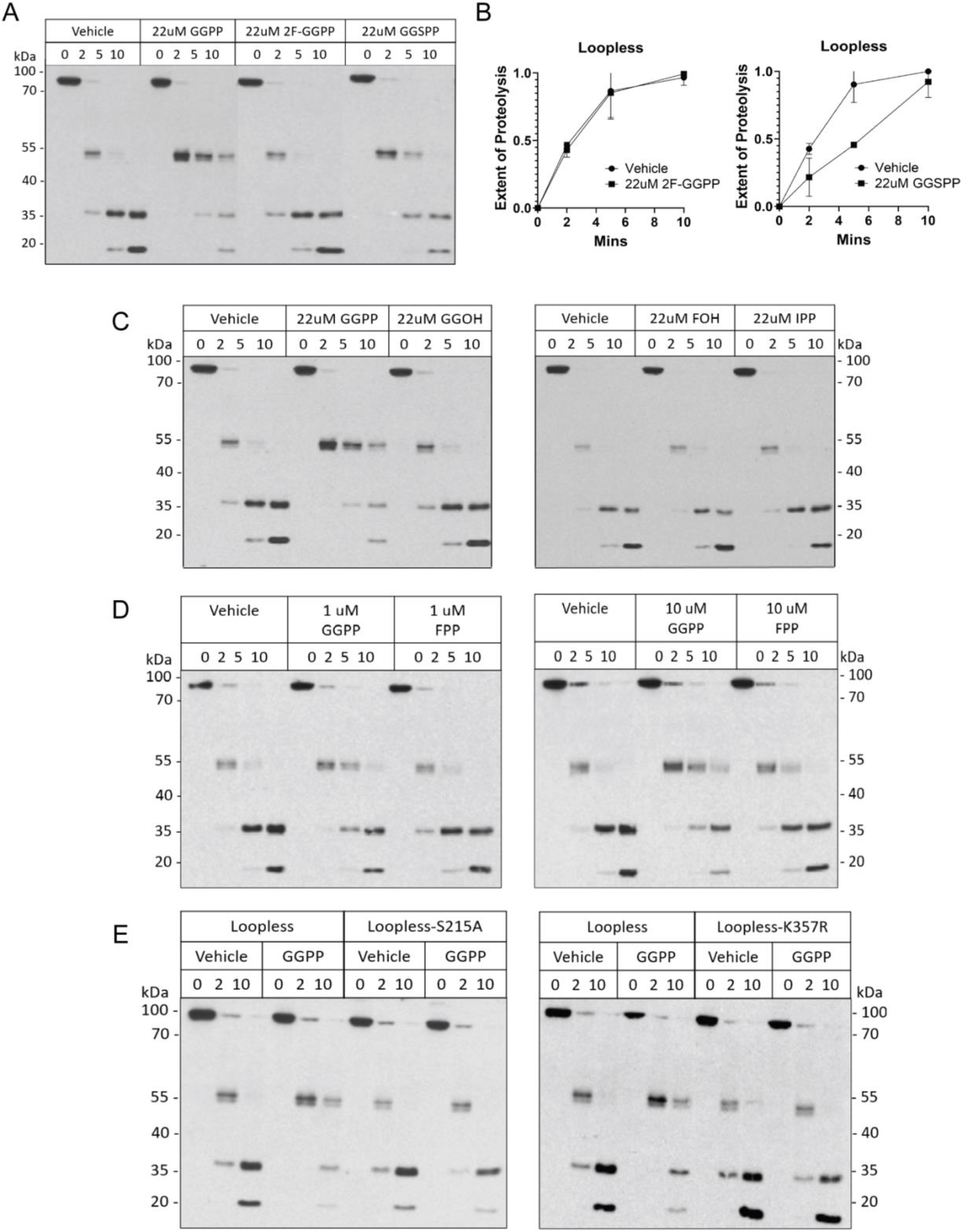
Loopless has the same ligand specificity as WT Hmg2 *in vitro*. *A*, 2F-GGPP has no effect while GGSPP has a stabilizing effect on the Loopless structure. Limited proteolysis of Loopless when treated with GGPP and compared to GGPP analogs 2F-GGPP and GGSPP over the indicated times. Note that GGSPP acted as an antagonist to GGPP with WT Hmg2’s structure. *B*, 2F-GGPP has no effect on the rate of proteolysis while GGSPP slows the rate of proteolysis. Extent of proteolysis was calculated for the effects of 2F-GGPP (not significant) and GGSPP (p < 0.05) on the Loopless structure. Loopless Extent of Proteolysis = (Doublet intensity/Total intensity). Values shown are means of three experiments. Error bars are SEM. Statistical significance was determined by two-way repeated-measures ANOVA followed by Sidak’s multiple comparisons test comparing Vehicle and GGPP at each time point. *C*, The early sterol pathway intermediates GGOH, FOH, and IPP do not have an effect on the Loopless structure. Limited proteolysis of the effect of the relevant GGPP early sterol pathway intermediates GGOH, FOH, and IPP on the Loopless structure over the indicated times. *D*, Loopless responds to low doses of GGPP and starts to respond at high FPP doses. Dose response of GGPP and FPP in the limited proteolysis assay at 1uM and 10uM, respectively. Loopless has a slight response to increasing doses of FPP but are not as significant as GGPP at the same nM doses. *E*, Loopless response to GGPP is SSD dependent. S215A and K357R are previously established SSD mutants that render WT Hmg2 unresponsive to GGPP in the limited proteolysis assay.

The striking similarity of interactions with GGPP analogs indicated that the binding site remaining in Loopless had very similar properties to that in full length Hmg2. We further tested this similarity by evaluating a number of mevalonate-derived isoprenoids that have no effect on the Hmg2 limited proteolysis assay. These include GGOH, FOH, and IPP (Fig. 5C). As with WT Hmg2-GFP, these molecules had no effect on the rate of Loopless proteolysis. In our extensive analysis of Hmg2 we learned that the 15-carbon isoprenoid FPP had a much weaker interaction with Hmg2 in the limited proteolysis assay, with a micromolar EC50 rather than the nanomolar potency reported for GGPP (12). Loopless retains the same nM level of specificity to GGPP and uM response to FPP in its in vitro response to these isoprenoids (Fig. 5D).

Finally, we have previously shown that GGPP-mediated Hmg2 response is dependent on a functional SSD. In addition to the S215A mutation, we have previously shown another example of a stable SSD mutation called K357R. The S215A mutation and the K357R mutation each rendered Loopless unresponsive to GGPP (Fig. 5E).

These data indicate that, when compared to Hmg2, the Loopless response to GGPP retains the same features in terms of GGPP structural specificity, potency, and SSD requirement. We next tested the ability of Loopless to multimerize, both to better understand this intriguing mutant, and to test its ability to participate in interactions with Hmg2 through trans effects such as toxic subunit interactions we discovered in our characterization of Hmg2 mallostery (24).

### Multimerization is preserved in the transmembrane domain

We have previously shown that Hmg2 forms homomultimers by direct biochemical means, and through observation of trans effects in which co-expressed mutants alter the GGPP response of wild-type Hmg2 (12). These studies were done with wild-type and full-length point mutants of wild-type Hmg2 containing both the L1L7 regions and the transmembrane region preserved in Loopless. To determine if the lumenal domain is required for Hmg2 multimerization, we co-expressed different WT Hmg2 and Loopless combinations with a GFP tag or a MYC tag for co-immunoprecipitation (Co-IP) experiments (Fig. 6A). The basic strategy is, after ensuring that co-expression of each protein is similar in lysates (“input”), the lysates are subject to immunoprecipitation with anti-GFP antibody, followed by immunoblot for both immunoprecipitation of the “bait” (GFP protein) and co-immunoprecipitation of the “prey” (MYC tagged protein) (Fig. 6B bottom). These data show that Hmg2 multimerizes with Loopless and Loopless can multimerize with other Loopless subunits. This suggests that multimerization is brought about by interactions within the transmembrane domain of Hmg2, since the absence of L1-L7 in either or both of the test proteins still allows strong immunoprecipitation signal. We next tested if co-expression of Loopless and Hmg2 caused any trans effects on GGPP response.

**Figure 6.**
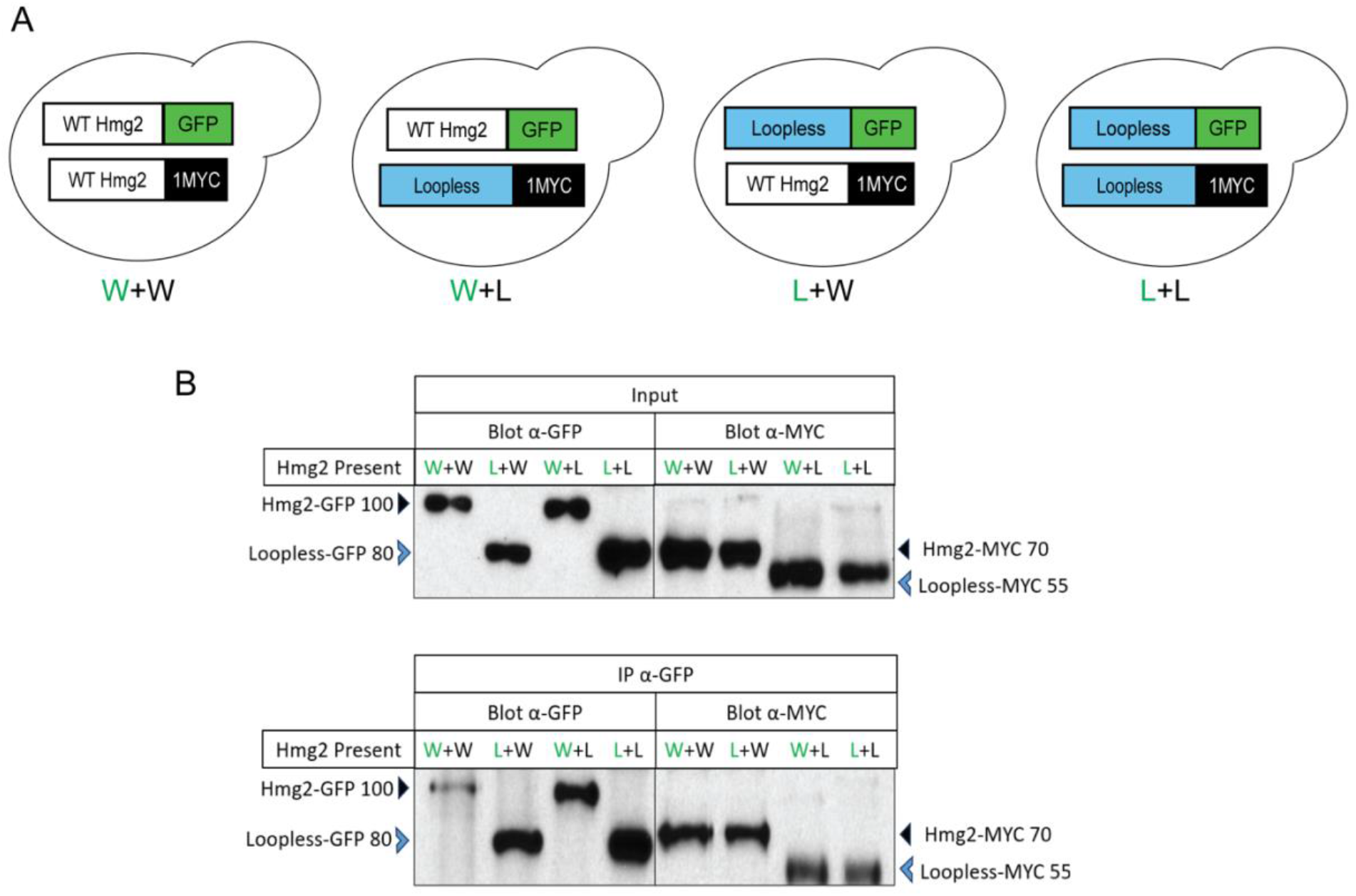
Loopless forms multimers with WT Hmg2 and other Loopless subunits. *A*, cartoon depiction of how yeast strains were constructed to express different Hmg2 or Loopless subunits. WT Hmg2 is depicted in a white box, Loopless is depicted in a blue box, GFP is depicted in a green box, and the 1 MYC tag is depicted in a black box. Strains are also denoted with a “W” for WT and “L” for Loopless as well as green font to represent which subunit is GFP-tagged and black font to represent which subunit is MYC-tagged. *B*, WT Hmg2 and Loopless subunits expressed in trans are detected and can be pulled down together after a GFP IP. western blots of Hmg2 or Loopless subunits isolated and solubilized with non-denaturing detergent from microsomes from co-expressed strains depicted in panel *A* blotted for GFP or MYC. Solubilized proteins are blotted for GFP or MYC as input for GFP immunoprecipitation (IP) (top). Solubilized proteins are IP’ed with a GFP antibody and blotted for GFP and MYC. Solubilized proteins are Hmg2 subunits are denoted with a filled black triangle and Loopless subunits are denoted with a blue open arrowhead.

### The lumenal domain is required for transducing a GGPP-mediated mallosteric response

Because Loopless forms multimers with WT Hmg2 and Loopless subunits, we wanted to test for any trans effects caused by their co-expression. We have previously shown that the co-expression of the highly stable Hmg2 S215A mutant with WT Hmg2 has a strong stabilizing effect on WT Hmg2 when treated with GGPP *in vivo* and *in vitro* (24).

First, we used a cycloheximide chase to measure the degradation kinetics of the co-expressed, fluorescent WT Hmg2-GFP with non-optical Loopless-MYC and compared it to WT Hmg2-GFP co-expressed with either Hmg2 S215A-MYC or WT Hmg2-MYC (Fig. 7A). As previously shown, co-expression of the highly stable, non-optical Hmg2-myc SSD mutant strongly blunted the GGPP response of co-expressed wild-type Hmg2-GFP (Fig. 7C)(23). Conversely, co-expressed non-optical Loopless had no significant effect on the degradation of WT Hmg2-GFP. We also compared the effects of the co-expression of Loopless-S215A-MYC on WT Hmg2-GFP and again found no significant trans effects on the degradation kinetics of WT Hmg2. Similarly, an optical Loopless-GFP was co-expressed with a non-optical WT Hmg2-MYC and compared to the co-expression of Loopless-GFP with Loopless-MYC. Again, Hmg2-myc had no effect on the highly stable Loopless-GFP when present in co-expression (Fig. 7B).

**Figure 7.**
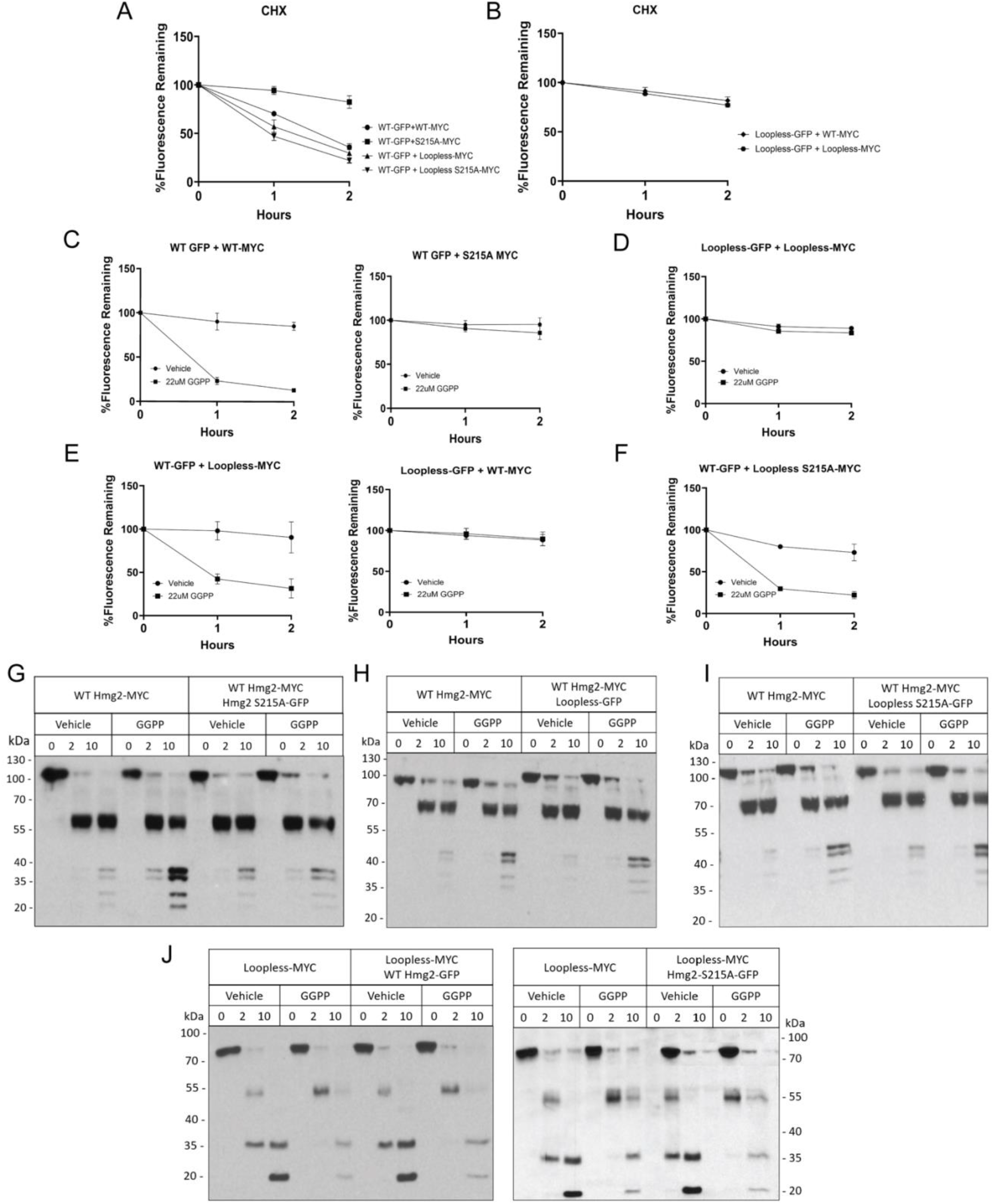
WT Hmg2 and Loopless show no trans effects *in vivo* or *in vitro*. All flow cytometry experiments show mean fluorescence normalized to time 0, from three separate cultures by flow cytometry of 10,000 cells each. Error bars are SEM. *A*, co-expression of dark (nonfluorescent) MYC-tagged, non-mallosteric mutant S215A strongly stabilizes WT Hmg2-GFP while co-expression of dark WT Hmg2-MYC, dark MYC-tagged Loopless, dark MYC-tagged Loopless S215A has no effect on WT Hmg2-GFP. WT Hmg2-GFP was co-expressed with dark WT Hmg2-MYC (filled circles), dark S215A Hmg2-MYC (filled squares), dark Loopless-MYC (filled up-triangle), and dark Loopless S215A-MYC (filled down-triangle). In a cycloheximide chase, dark S215A Hmg2-MYC slows the degradation rate of WT Hmg2-GFP relative to the strains that co-express dark WT Hmg2-MYC, dark Loopless-MYC, or dark Loopless-S215A-MYC, respectively. *B*, co-expression of dark MYC-tagged WT Hmg2 or dark MYC-tagged Loopless has no effect on the degradation rate of Loopless-GFP. Loopless-GFP was co-expressed with dark WT Hmg2-MYC (filled diamond) or dark Loopless-MYC (filled hexagon). In a cycloheximide chase, there are no changes to the degradation rate when Loopless-GFP is co-expressed with dark WT Hmg2-MYC or dark Loopless-MYC. *C*, co-expression of dark S215A Hmg2-MYC inhibited the response of WT Hmg2-GFP to GGPP when compared to the co-expression of dark WT Hmg2-MYC. Co-expression of an Hmg2 mutant with insensitivity to GGPP reduces GGPP-induced degradation of WT Hmg2-GFP. Filled circles show vehicle control and filled squares show GGPP treatment. *D*, co-expression of dark Loopless-MYC does not have an effect on Loopless-GFP response to GGPP *in vivo*. *E*, co-expression of dark Loopless-MYC does not affect WT Hmg2-GFP response to GGPP and dark WT Hmg2-MYC does not affect Loopless-GFP response to GGPP. WT Hmg2-GFP co-expressed with a mutant that does not undergo GGPP-induced degradation still degrades over time when treated with GGPP (left). Loopless-GFP co-expressed with WT Hmg2-MYC which does undergo GGPP-induced degradation remains stable over the indicated time (right). *F*, co-expression of dark Loopless-S215A mutant does not have a stabilizing effect on GGPP-induced degradation of WT Hmg2-GFP. WT Hmg2-GFP co-expressed with the dark SSD mutant Loopless-S215A still responds and undergoes GGPP-induced degradation. *G*, co-expression of a non-mallosteric SSD Hmg2 mutant strongly blocked GGPP-induced misfolding of WT Hmg2-MYC *in vitro*. GGPP-induced misfolding increased proteolytic cleavage for WT Hmg2 myc but is attenuated when the Hmg2 S215A-GFP mutant is co-expressed. *H*, co-expression of Loopless-GFP does not affect the WT Hmg2-MYC GGPP-induced misfolding *in vitro*. WT Hmg2-MYC expressed with Loopless GFP did not affect the rate of proteolysis in a limited proteolysis assay when GGPP is added. *I*, co-expression of the SSD mutant Loopless-S215A-GFP does not affect the WT Hmg2-MYC GGPP-induced misfolding *in* vitro. WT Hmg2-MYC expressed with the SSD mutant Loopless-S215A-GFP did not affect the rate of proteolysis in a limited proteolysis assay when GGPP is added. J, co-expression of WT Hmg2-GFP or a non-mallosteric SSD mutant WT Hmg2-S215A-GFP does not change the Loopless-MYC stabilizing response to GGPP *in vitro*. Loopless-MYC expressed with WT Hmg2-GFP (left) or a non-mallosteric SSD mutant Hmg2 S215A-GFP (right) do not change the Loopless-MYC rate of proteolysis when GGPP is added.

GGPP effects in these co-expression strains were also tested by examining GGPP-dependent alteration in steady state levels as above. As previously reported (24), co-expression of the stable SSD mutant Hmg2 S215A has a stabilizing effect on WT Hmg2 when compared to the co-expression of an optical WT Hmg2 with a non-optical WT Hmg2 (Fig. 7C). The co-expression of Loopless subunits had no effect on the *in vivo* changes in Hmg2-GFP, nor were Loopless-GFP levels altered by GGPP when Hmg2 was co-expressed (Fig. 7D). When WT Hmg2-GFP is co-expressed with Loopless-MYC or when Loopless-GFP is co-expressed with WT Hmg2-MYC, there is no effect on their respective changes in steady state levels when GGPP is added (Fig. 7E). Similarly, there is no change in steady state levels when WT Hmg2-GFP is co-expressed with Loopless-S215A-MYC and treated with GGPP (Fig. 7F). Thus, despite retention of the ability for the stable Loopless mutant to interact with Hmg2 as multimers, there were no observable *in vivo* trans interactions: Hmg2 did not impart regulation to co-expressed Loopless-GFP, nor did Loopless alter Hmg2-GFP’s response to GGPP.

We also analyzed these mutants for possible trans effects with the limited proteolysis assay. This is the case with full-length non-responsive Hmg2 mutants: co-expression of Hmg2 S215A-GFP strongly blunts the effects of GGPP on WT Hmg2-MYC (Fig. 7G). In contrast, when WT Hmg2-MYC is co-expressed with Loopless-GFP, there was no detectable change in the GGPP effect on Hmg2-MYC proteolysis when compared to WT Hmg2-MYC alone (Fig. 7H). Similarly, co-expression of Loopless-S215A-GFP had no effect on the GGPP response of Hmg2-MYC (Fig. 7I). And furthermore, neither co-expression of WT Hmg2-GFP, nor S215A-Hmg2-GFP had any effect on the GGPP response of Loopless-MYC when compared to Loopless-MYC alone (Fig. 7J).

Together, these data suggest that while Hmg2 and Loopless form multimers, there is no mallosteric communication between these subunits, as assessed by co-expression experiments, and in striking contrast to the observed trans effects using various combinations of full-length Hmg2. This suggests that the lumenal domains are required for the mallosteric response of Hmg2, and for communication between subunits that is a hallmark of classic allostery and mallostery.

## Discussion

In this work, we sought to investigate the nature and location of the Hmg2 GGPP-binding site to further our understanding of GGPP-mediated Hmg2 mallostery. Together with the AlphaFold model of Hmg2 an analogy to Scap, we initially hypothesized that the GGPP-binding site was in Hmg2’s lumenal domain. A saturated single point mutation search in the lumenal domain did result in a collection of mutants that were more stable than WT Hmg2, but in contrast to the SSD S215A mutant isolated long ago, none completely removed response *in vitro*. Certain loop 7 mutants also yielded an interesting pharmacological chaperoning response to GGPP; meaning that when GGPP was added, the limited proteolysis assay revealed a ligand-dependent slowing of proteolysis, as opposed to the mallosteric hastening seen in wild-type Hmg2p. This screen clearly shows that the native structure of Loops 1 and 7 are critical for allowing GGPP-dependent mallostery that causes enhancement of Hmg2 entry into HRD quality control pathway. Furthermore, our analysis of the Loopless and L1L7 constructs imply, in contrast to the abundantly tested Scap model, that the Hmg2 transmembrane domain contains a GGPP-binding site. Detailed testing of the GGPP effects on Loopless showed that this binding site has all the features previously observed in WT Hmg2: structural specificity of the GGPP structure, ability for only the GGSPP molecule to bind, and low potency of the FPP molecule. Furthermore, the Loopless – GGPP interaction required a functional SSD, while the mallosteric response requires the presence of an intact lumenal domain (Fig. 8). The SSD has long been considered a region where sterols like cholesterol interact and trigger the protein’s conformational change, but not necessarily the region of ligand-binding like in the case of Scap. Recent work has further strengthened this conceptual link and shows that ligand binding to mammalian HMGR induces conformational changes required for its regulated degradation, highlighting a greater degree of conservation with yeast Hmg2 than previously understood (29). Together, these findings support a revised model in which GGPP binding occurs within the transmembrane domain, while the lumenal domain is required to transduce this binding event into a mallosteric degradation signal.

**Figure 8.**
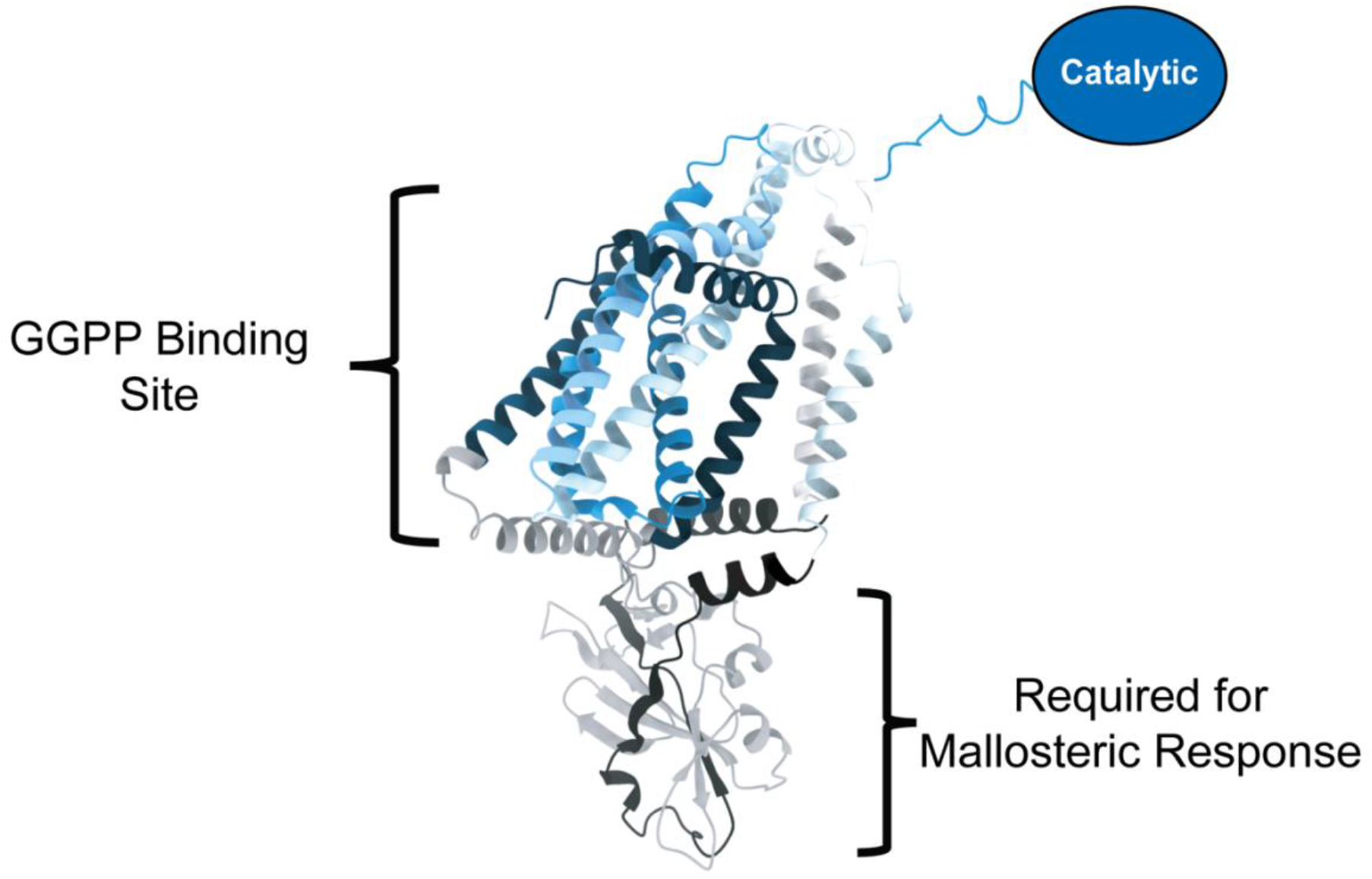
A new model for GGPP-mediated Hmg2 Mallostery. A cartoon schematic of the AlphaFold model of Hmg2 that highlights the function of the transmembrane domain and lumenal domain. The GGPP binding site is retained in the Hmg2 transmembrane domain (top bracket). The lumenal domain is required to transduce a GGPP-mediated mallosteric response (bottom bracket).

Importantly, the response to GGPP by Loopless is not the wild-type “mallosteric” response hallmarked by increased proteolytic sensitivity in vitro and hastened HRD pathway entry in vivo. Instead, the interaction of GGPP with Loopless has the features of a pharmacological chaperone, in which the structure appears to become more compact and thus harder to proteolyze. The many examples of pharmacological chaperones indicate that the more typical response of a misfolded, ligand-binding protein is to show improved folding when ligand is present. Interestingly, in other work, we have explored the mutagenic acquisition of ligand-dependent stability effects on a model protein to explore pharmacological chaperoning vs mallostery. We used stable, GFP-tagged chorismate mutase (CM-GFP), which undergoes allosteric regulation by tryptophan binding. Using similar saturation methods as in the studies herein, we generated a large collection of CM-GFP point mutants with varying stabilities. We looked for two classes of mutants: CM variants that underwent degradation that was stabilized by the presence of tryptophan, and we looked for mutants in which tryptophan caused increased destabilization. That is, we looked for point mutants that caused normally stable CM to respond to tryptophan as a pharmacological chaperone, or, CM mutants that would respond to tryptophan as a mallosteric ligand. Remarkably, we obtained a variety of destabilized CM mutants that were stabilized by Trp binding to its allosteric site. In striking contrast, we have so far never found a CM mutant that becomes less stable in the presence of its ligand, despite hundreds of person-hours of diligent screening (30). Thus, it appears that pharmacological chaperoning is acquired more easily, in terms of mutagenic search, than mallostery. So, it is perhaps not surprising that the GGPP binding site of Loopless can bring about a pharmacological chaperone effect when the information of the lumenal loops is absent.

Discovery of the Loopless mutant now leads to several open questions. Loopless was still able to form multimers with Hmg2 subunits and to itself, suggesting that the site of multimerization is retained in the transmembrane domain. Would disrupting the site of multimerization affect Hmg2’s regulation and response to GGPP? Future studies combining saturation mutagenesis of the transmembrane domain with in vitro structural assays may help identify residues directly involved in GGPP binding and further refine the location of the interaction site. Clearly what is now needed for understanding this remarkable regulatory axis is a detailed structure of Hmg2 with and without the GGPP ligand. We have previously shown that solubilized Hmg2 responds to GGPP (24), and thus the state is set for attainment of a physiologically meaningful structure.

This study leveraged the use of the AlphaFold model of Hmg2 to ask hypothesis-driven questions about the relationship between GGPP and the structure of Hmg2. While AlphaFold has become the most successful publicly available tool for predicting protein structure, the current algorithm does not always accurately predict unsolved protein structures. This tends to be the case especially for membrane proteins because AlphaFold is not consistent with its placement of transmembrane domains (31). AlphaFold was a useful tool for further interrogating the structure of Hmg2 by helping us design the Loopless mutant. Ultimately, we will better understand Hmg2 GGPP-mediated mallostery by solving the structure of Hmg2 with more direct structural techniques.

The field of targeted protein degradation has largely focused on the engineering of bivalent degraders called proteolysis targeting chimeras (PROTACs) that are structured to have tunable target binding attached to a linker that recruits a ubiquitin ligase for marking the target for degradation (32). While PROTACs theoretically have the therapeutic potential to target specific disease-related proteins, they also have their limitations. PROTACs are relatively large and are not readily cell-permeable, making tissue-specific delivery a current challenge (33). In the case of PROTACs as a cancer therapeutic, current PROTACs designs recruit E3 ligases that have been altered in numerous cancer types, creating the need to expand PROTACs-mediated degradation to other E3 ligases. Clinical treatment with PROTACs is also still early and requires more observations of off-target and downstream effects.

Despite this emphasis on engineered bivalent degraders, endogenous monovalent mechanisms, meaning molecules that only bind to the target (such as Hmg2) but trigger degradation, remain poorly understood. Our work on studying GGPP-mediated Hmg2 mallostery provides a well-studied example of a natural monovalent degrader, a class of small molecules that promote the degradation of specific proteins. In this system, the endogenous sterol pathway intermediate GGPP binds Hmg2 and induces a subtle, regulated misfolding event that is sufficient to engage ER-associated degradation without requiring direct recruitment of ubiquitin ligases. As many therapeutically relevant proteins are governed by conformational stability rather than catalytic activity, understanding GGPP-mediated Hmg2 misfolding provides a powerful conceptual framework for expanding monovalent degrader strategies with translational potential.

## Methods and Materials

### Reagents

GGPP, GGOH, FPP, FOH, IPP, cycloheximide, trypsin, and digitonin were purchased from Sigma-Aldrich. Lovastatin was a gift from Merck & Co (Rahway, NJ). GGSPP and 2-fluoro-GGPP were gifts from Reuben Peters (Iowa State University) and Philip Zerbe (University of California, Davis). Lyophilized GGSPP was also purchased from Echelon Biosciences. Anti-myc 9E10 supernatant was prepared from cells (CRL 1729, American Type Culture Collection) cultured in DMEM medium (GIBCO BRL) with 10% fetal calf serum. Living colors mouse anti-GFP monoclonal antibody was purchased from Clontech, and HRP-conjugated goat anti-mouse antibody was purchased from Jackson ImmunoResearch.

### Yeast strains and plasmids

Plasmids (Table S1) and yeast strains (Table S2) were constructed by standard techniques. Yeast strains were isogenic and derived from the S288C background. Yeast strains were grown in minimal media (Difco yeast nitrogen base supplemented with necessary amino and nucleic acids) with 2% glucose or rich medium (YPD). Strains were grown at 30 °C with aeration. Constructs were introduced by integration of plasmid cut with StuI at the ura3-52 locus, BsgI at the trp1 locus, or with PpuMI at the leuΔ promoter.

### Saturation Mutagenesis Library Preparation

Single-stranded 150bp oligos containing single-point amino acid mutations to indicated regions of HMG2 were synthesized by IDT. Single stranded oligos were extended with standard PCR techniques to form double-stranded products for insertion into plasmid backbone by standard Gibson cloning techniques. Each library was analyzed by deep sequencing methods to determine coverage of the library. Libraries were analyzed in RStudio. Libraries were then transformed into a yeast strain with a BTS1 overexpression construct for constitutive high levels of GGPP.

### Saturation Mutagenesis Screening

Stably fluorescing yeast colonies detected under the 488nm light with a wratten lens were colony purified by streaking onto fresh dropout plates. Mutant Hmg2 plasmids were purified by growing colony purified candidates in YPD overnight and extracted with standard yeast plasmid purification techniques. Plasmids were sequenced and re-transformed into an S288C background with normal sterol flux for *in vivo* and *in vitro* analysis.

### Flow cytometry

Flow cytometry experiments were performed as previously described(12). Yeast cultures were grown in minimal media to early log phase. Indicated molecules (22 μM GGPP or 50 μg/ml cycloheximide) or equal volumes of vehicle (7:3 methanol:10 mM ammonium bicarbonate for GGPP) were added directly to culture medium. Cell fluorescence was measured for 10,000 cells per condition using a BD Accuri C6 Plus flow cytometer (BD Biosciences). Any averages shown are means of 10,000 ungated events.

### Fluorescence Microscopy

Yeast strains expressing WT Hmg2-GFP, S215A-GFP, Y48D-GFP, T424P-GFP, I479D-GFP, L1L7-sfGFP, and Loopless-GFP were grown to early log phase in minimal media. 2 µL of concentrated cells were spotted onto a glass slide and covered with a coverslip for live-cell imaging. Cells were imaged at 100× magnification with oil immersion on a Carl Zeiss Axioskop 2 MOT microscope (Carl Zeiss Microscopy, Göttingen, Germany) using a 488 nm laser for GFP excitation. Images were captured with an AxioCam HRm digital camera (Carl Zeiss MicroImaging GmbH, Göttingen, Germany) and processed using ImageJ (NIH).

### Microsome preparation

Microsomes were prepared as described previously(12). Yeast strains were grown to midlog phase in YPD. 10 OD eq were resuspended in 240 uL of lysis buffer (0.24 M sorbitol, 1mM EDTA, 20 mM KH2PO4/K2HPO4, pH 7.5) with protease inhibitors (2 mM phenylmethylsulfonyl fluoride and 142 mM tosyl-phenylalanyl chloromethyl ketone). Acid-washed glass beads were added to the meniscus, and cells were lysed at 4 °C on a multivortexer for six 1-min intervals with 1 min on ice in between. Lysates were cleared by centrifugation in 5-s pulses until no pellet was apparent. Microsomes were pelleted from cleared lysates by centrifugation at 14,000 x g for 5 min, washed once in XL buffer (1.2 M sorbitol, 5 mM EDTA, 0.1 M KH2PO4/K2HPO4, pH 7.5), and resuspended in XL buffer.

### Limited proteolysis assay

Microsomes were subjected to limited proteolysis as described previously(12). Briefly, resuspended microsomes were treated with the indicated isoprenoid molecules or equal volumes of vehicle and then incubated with trypsin at a final concentration of 15ug/ml at 30 °C. Reactions were quenched at the times indicated with an equal volume of 2x urea sample buffer (USB; 8 M urea, 4% SDS, 1 mM DTT, 125 mM Tris base, pH 6.8). Samples were resolved by 14% SDS-PAGE, transferred to nitrocellulose in 15% methanol, and blotted with 9E10 anti-myc antibody.

### Detergent Solubilized Limited Proteolysis

Detergent solubilized microsome subject to limited proteolysis as described previously (24). The limited proteolysis assay was performed as described above with the following variations: After initial pelleting and washing of microsomes, microsomes were first thoroughly resuspended in XL buffer and then solubilized by the addition of detergent solution at 10x the desired final concentration in XL buffer 1% digitonin. Preparations with detergent were incubated at 4C for 1 h with rocking and then repeatedly pipetted up and down. Finally, samples were cleared by centrifugation in a benchtop microcentrifuge for 15 min at 16,000g. The supernatants were then separated by ultracentrifugation at 89,000 RPM for 15 min, and the supernatant from this step was used for the proteolytic assay. This assay was identical to the limited proteolysis assay described above, except that a lower concentration of trypsin (3 μg/ml instead of 15) was used.

### Coimmunoprecipitation of Hmg2 Subunits

Coimmunoprecipitation of Hmg2 subunits was performed as described previously (24). Microsomes in IPB were incubated at 4 °C for 1 h with rocking. Microsomes were pipetted up and down repeatedly, and then solutions were cleared by centrifugation at 14,000xg for 15 min. Supernatants were incubated with 5 ul of monoclonal mouse anti-GFP antibody overnight at 4 °C with rocking. After overnight incubation, 100 ul of a 50% protein A–Sepharose bead slurry swelled in IP buffer without detergent were added. Samples were incubated at 4 °C for 2 h with rocking. Beads were then pelleted for 30 s at low speed and 1 min by gravity and washed twice with IPB and once with IP wash buffer (100 mM NaCl, 10mM Tris base, pH 7.5). Beads were aspirated to dryness and resuspended in 2x USB. Samples were resolved by electrophoresis on 10% polyacrylamide gels, transferred to nitrocellulose in 12% methanol buffer, and immunoblotted with anti-GFP or anti-myc antibody as indicated.

