## Supplemental Figure 1 for "Structural Determinants of Ligand-Mediated Hmg2 Mallostery"

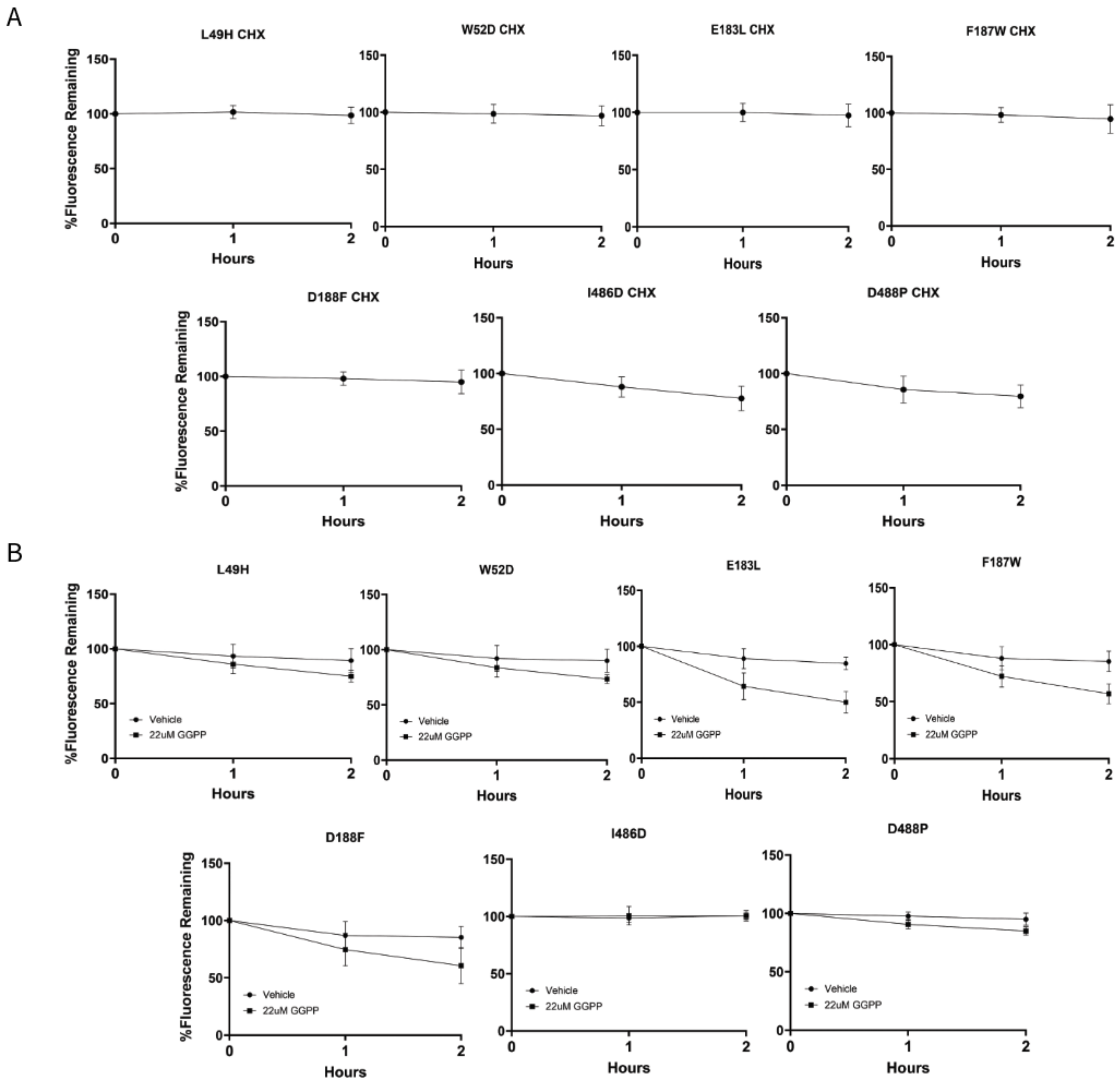


**Supplemental Figure 1. Other Loop 1 and Loop 7 single point mutants that are non-responsive to GGPP.**

All flow cytometry experiments show mean fluorescence normalized to time 0, from three separate cultures by flow cytometry of 10,000 cells each. Error bars are SEM. *A*, L49H. W52D, E183L, F187W, and D188F are Loop 1 mutants. I486D and D488P are Loop 7 mutants. Loop 1 and Loop 7 mutants were stable compared to the non-responsive SSD mutant S215A. The degradation rate of Hmg2-GFP, SSD mutant S215A, Loop 1, and Loop 7 mutants over the indicated times after addition of cycloheximide (CHX). *B*, Loop 1 and Loop 7 mutants have a range of sensitivity to 22uM of GGPP. The effect of GGPP on Hmg2-GFP and Hmg2 mutants loop mutants on steady-state levels over the indicated times.
