## Supplemental Table 1 for "Structural Determinants of Ligand-Mediated Hmg2 Mallostery"

**Supplemental Table 1. Plasmids used in this work**

| Plasmid | Genotype |
|---|---|
| pRH469 | URA3::TDH3pr-HMG2-GFP YIp |
| pRH680 | LEU2::TDH3pr-HMG2-GFP YIp |
| pRH1581 | URA3::TDH3pr-1myc$_L$-HMG2-GFP YIp |
| pRH1693 | URA3::TDH3pr-K357R-1myc$_L$-HMG2-GFP YIp |
| pRH2177 | URA3::TDH3pr-S215A-1myc$_L$-HMG2-GFP YIp |
| pRH2250 | URA3::TDH3pr-HMG2-1myc YIp |
| pRH3211 | LEU2::TDH3pr-S215A-HMG2-GFP YIp |
| pRH3212 | URA3::TDH3pr-S215A-HMG2-1myc YIp |
| pRH3231 | URA3::TDH3pr-HMG2-1myc-GFP YCp |
| pRH3287 | URA3::TDH3pr-L1L7-1myc-sfGFP YIp |
| pRH3312 | URA3::TDH3pr-HMG2-Loopless-1myc-1flag-GFP |
| pRH3345 | URA3::TDH3pr-1myc$_L$-HMG2 YIp |
| pRH3347 | TRP1:: TDH3pr-HMG2-Loopless-1myc-1flag-GFP |
| pRH3348 | URA3:: TDH3pr-HMG2-Loopless-1flag-GFP |
| pRH3349 | TRP1:: TDH3pr-HMG2-Loopless-1myc-1flag |
| pRH3350 | URA3:: TDH3pr-S215A-HMG2-Loopless-1myc-1flag-GFP |
| pRH3351 | URA3:: TDH3pr-K357R-HMG2-Loopless-1myc-1flag-GFP |
| pRH3352 | URA3: TDH3pr-S215A-HMG2-Loopless-1flag-GFP |
| pRH3353 | URA3: TDH3pr-S215A-HMG2-Loopless-1myc-1flag |
| pRH3354 | URA3:: TDH3pr-HMG2-Loopless-1flag-1myc |
