## Supplemental Table 2 for "Structural Determinants of Ligand-Mediated Hmg2 Mallostery"

**Supplemental Table 2. Yeast strains used in this work**

| Strain | Genotype | Reference |
|---|---|---|
| RHY7661 | MAT*a ade2-101 met2 his3Δ200 lys2-801 leu2Δ ura3-52::URA3::TDH3pr-1myc*$_L$*-HMG2-GFP trp1::hisG* | Garza et al. 2009 |
| RHY7683 | MAT*a ade2-101 met2 his3Δ200 lys2-801 leu2Δ ura3-52::URA3::TDH3pr-S215A-1myc*$_L$*-HMG2-GFP trp1::hisG* | Theesfeld et al. 2011 |
| RHY11115 | MAT*a ade2-101 met2 his3Δ200 lys2-801 leu2Δ::LEU2::TDH3pr-Hmg2-GFP ura3-52::URA3::TDH3pr-HMG2-1myc trp1::hisG* | Wangeline & Hampton, 2021 |
| RHY11128 | MAT*a ade2-101 met2 his3Δ200 lys2-801 leu2Δ ura3-52::URA3::TDH3pr-K357R-1myc*$_L$*-HMG2-GFP trp1::hisG* | Wangeline & Hampton, 2018 |
| RHY12258 | MAT*a ade2-101 met2 his3Δ200 lys2-801 leu2Δ::TDH3pr-BTS1 ura3-52 trp1::hisG* | This work |
| RHY12742 | MAT*a ade2-101 met2 his3Δ200 lys2-801 leu2Δ ura3-52::URA3::TDH3pr-L1L7-1myc-GFP trp1::hisG* | This work |
| RHY12759 | MAT*a ade2-101 met2 his3Δ200 lys2-801 leu2Δ ura3-52::URA3::TDH3pr-HMG2-Loopless-1myc-1flag-GFP trp1::hisG* | This work |
| RHY12833 | MAT*a ade2-101 met2 his3Δ200 lys2-801 leu2Δ::LEU2::TDH3pr-Hmg2-GFP ura3-52::URA3::TDH3pr-HMG2-Loopless-1flag-1myc trp1::hisG* | This work |
| RHY12834 | MAT*a ade2-101 met2 his3Δ200 lys2-801 leu2Δ ura3-52::URA3::TDH3pr-HMG2-1myc trp1::hisG::TRP1 TDH3pr-HMG2-Loopless-1flag-GFP* | This work |
| RHY12835 | MAT*a ade2-101 met2 his3Δ200 lys2-801 leu2Δ ura3-52::URA3:: TDH3pr-HMG2-Loopless-1flag-1myc trp1::hisG::TRP1 TDH3pr-HMG2-Loopless-1flag-GFP* | This work |
| RHY12836 | MAT*a ade2-101 met2 his3Δ200 lys2-801 leu2Δ::LEU2::TDH3pr-Hmg2-GFP ura3-52::URA3::TDH3pr-S215A-HMG2-Loopless-1flag-1myc trp1::hisG* | This work |
| RHY12837 | MAT*a ade2-101 met2 his3Δ200 lys2-801 leu2Δ ura3-52::URA3::TDH3pr-Hmg2-1myc trp1::hisG::TRP1 TDH3pr-S215A-HMG2-Loopless-1flag-GFP* | This work |

| | | |
|---|---|---|
| RHY12838 | MAT*a ade2-101 met2 his3Δ200 lys2-801 leu2Δ ura3-52::URA3::TDH3pr-S215A-Hmg2-GFP trp1::hisG::TRP1 TDH3pr-HMG2-Loopless-1flag-1myc* | This work |
